# Topical siRNA therapy of diabetic-like wound healing

**DOI:** 10.1101/2023.11.06.565704

**Authors:** Eva Neuhoferova, Marek Kindermann, Matej Buzgo, Karolina Vocetkova, Dalibor Panek, Petr Cigler, Veronika Benson

## Abstract

Non-healing wounds are a serious complication in diabetic patients. One of the detrimental factors contributing to limited wound healing is the accumulation of metalloproteinase-9 (MMP-9) in the wound. Selective inhibition of MMP-9 is one of the established therapeutic targets for diabetic wound healing and is therefore of great interest. Here we focused on development of gene silencing system for localized delivery of antisense siRNA against MMP-9 into the wound. We have developed a functional and biocompatible wound dressing allowing controlled release of a traceable vector loaded with the target siRNA. Specifically, the dressing consists of a degradable scaffold of polymer nanofibers embedded with the vector nanosystem, polymer-coated fluorescent nanodiamonds (FNDs). The biocompatible cationic polymer shell on FNDs was designed and optimized for binding of siRNA and formation of colloidally stable FND-siRNA complexes in physiological environment. A hybrid nanofiber scaffold consisting of poly(vinyl alcohol) and poly(caprolactone) ensured continuous release of FND-siRNA complexes from the dressing. The photostable luminescence of FNDs allowed us to monitor the vector system in the wound. Our dressing was tested on murine fibroblasts and also applied to wounds in a diabetic murine model to evaluate its suitability in terms of *in vivo* toxicity, biological efficacy and manipulation. The treatment resulted in significant local inhibition of MMP-9 and reduction of the wound healing time. The scar formation for treated diabetic-like animals became comparable with non-treated diabetes-free mice. Our results suggest that the application of our biocompatible dressing loaded with a non-toxic vector nanosystem is an effective and promising approach in gene therapy of non-healing wounds.

## 1. Introduction

Cutaneous wound healing is a physiological process involving a complex overlap of cascade events resulting in the reconstruction of the injured skin [1,2]. Once the wound does not follow the normal healing pattern (hemostasis, inflammation, proliferation and remodeling phase) within a period of roughly 6-8 weeks, the human wound is typically considered as a chronical one [3]. Due to the high complexity of the healing process, the chronic wounds are quite common [4]. Among numerous causes of impaired wound healing [5], diabetes mellitus attracts attention as a growing international health concern with no end in sight [3,6]. The chronicity (wound healing delay) observed in diabetes mellitus can be triggered via different mechanisms including prolonged inflammation, poor blood (oxygen) delivery, sustained infection, growth factors degradation, increased levels of matrix metalloproteinases (MMPs) and others [3,7]. Thus, targeting these pathological triggers can serve as an efficient tool for the faster treatment of chronic diabetic wounds, which represent a heavy burden on public health and the healthcare system [8].

Despite the above-mentioned findings in elucidating wound repair mechanisms, the most common treatment of non-healing diabetic ulcers is based on removal of non-viable tissue from the wound followed by conventional dressing – gauzes, semipermeable films, hydrocolloids, alginates etc. [4,9]. This kind of dressing has typically only limited availability to bring the chronicity causing triggers back to the normal levels and thus facilitating the wound to heal faster [4,10]. On the other hand, a lot of effort has been taken to develop bioactive and biodegradable dressings containing antimicrobial agents [11,12], materials for gene therapy [13–16] or even living cells [4,17,18]. These active compounds or cells can be effectively released from the biodegradable scaffold due to hydrolytic enzymes in the wound exudates [19], the presence of bacteria in the infected wound [20] or other, rather mechanical, effects [3]. Although these dressings could significantly reduce hospitalization time, their preparation is costly and they still have not been used widespread [4].

Recently, electrospinning technology applied for the fabrication of biodegradable/biomimetic nanofiber dressing has gained exceptional interest due to its simplicity, robustness, and possibility to construct even portable electrospinning machines [21] offering an attractive way for tailored wound care [2,9]. Nanofiber character of the dressing and its high porosity allows embedding of bioactive moieties, absorption of the wound exudate, wound hydration, and permeation of oxygen [22]. Furthermore, in contrast with conventional dressing, the nanofiber arrangement mimics the structure of the extracellular matrix (ECM) and thus promotes attachment, proliferation and migration of the cells in the wound [23].

A crucial presumption for successful wound reconstruction is an appropriate ECM remodeling (final phase of wound healing) and the formation of primary scar [1]. Thus, it is obvious that a balanced synthesis–degradation process of ECM is essential. Degradation of damaged ECM during wound healing is driven by the zinc-dependent MMPs. During the normal healing process (acute wound without comorbidities), the MMPs are transiently expressed and extracellularly activated to obtain their proteolytic activity [1,24]. However, persisting presence of elevated MMP levels and abnormally low levels of tissue inhibitors of metalloproteinases were found in chronic wounds [24,25]. This pathological state is leading to the undesirable imbalance of ECM synthesis–degradation process and ECM loss.

In this work, we present a biodegradable nanofiber dressing, which mediates downregulation of MMP-9 (gelatinase B) levels in wounds *via* RNA interference mechanism. Our approach combines advantages of the synthetic nanofiber mesh (NF) embedded with a non-toxic nanosystem delivering the siRNA targeted against MMP-9 mRNA in the cells. As delivery particles we use recently developed hybrid nanosystem based on nanodiamonds coated with a cationic statistical copolymer poly{(2-dimethylaminoethyl methacrylate)-*co*-[NL(2-hydroxypropyl) methacrylamide]} (Cop^+^-FND) [26,27]. The complex of Cop^+^-FND with siRNA, Cop^+^-FND:siRNA, eliminates enzymatic cleavage of siRNA, is non-toxic and is effective as *in vivo* delivery platform [28]. The nanofiber mesh composite NF{Cop^+^-FND:siRNA} is fabricated by electrospinning and has two functions: primary bioactive dressing and a scaffold for controlled release of Cop^+^-FND:siRNA complexes into the wound. In addition, the nanodiamonds contain nitrogen–vacancy defects in the crystal lattice which produce a near-infrared fluorescence upon green laser excitation and have relatively long emission lifetime (≈20–40 ns, [29]). We use these specific features to easily distinguish the nanodiamonds from the polymer scaffold autofluorescence [28]. Fluorescent lifetime imaging provides us an in depth information about the spatial distribution of Cop^+^-FND:siRNA complexes inside the nanofibers. Finally, we test the performance of NF{Cop^+^-FND:siRNA} system on murine fibroblasts and in a diabetic murine wound model to evaluate its suitability in terms of *in vitro* and *in vivo* toxicity, biological efficacy and manipulation. We evaluate and compare the inhibition of MMP-9, the wound healing time and the scar formation in treated diabetic-like and diabetes-free mice.

## 2. Materials and methods

### 2.1. Materials

NIH/3T3 CRL-1658™ cells (ATCC®) were obtained from the Institute of Experimental Medicine, CAS, Czech Republic. TaqMan Universal Master Mix, MMP-9 specific qPCR assay (Mm00442991_m1), High-Capacity cDNA Reverse Transcription Kit, Power SYBR™ Green PCR Master Mix, LDH Cytotoxicity Assay Kit, RNAlater^TM^ Stabilization Solution, M-PER™ Mammalian Protein Extraction Reagent, Pierce™ Coomassie Plus™ (Bradford) Assay Kit, Nunc™ MicroWell™ 96-Well Microplates, Novex™ Tris-Glycine SDS Sample Buffer (2X), Novex™ Sharp Pre-stained Protein Standard, SimplyBlue™ SafeStain, NuPAGE™ LDS Sample Buffer (4X), NuPAGE™ Sample Reducing Agent (10X), NuPAGE™ 4-12% Bis-Tris Mini gel, SuperSignal™ West Pico PLUS Chemiluminescent Substrate, and XCell SureLock Mini-Cell Electrophoresis System were purchased from Thermo Fisher Scientific. Dulbecco’s modified Eagle’s medium (DMEM), fetal bovine serum (FBS), Streptozotocin and duplexed siRNA against MMP-9 mRNA (MW = 14,604) sense strand: 5’-r(GCC UAU UUC UGC CAU GGC AAA) d(AA)-3’ and antisense strand: 5’-r(UUU GCC AUG GCA GAA AUA GGC) d(TT)-3’ were purchased from Sigma–Aldrich. Fluorescently labelled siRNA duplex against GAPDH mRNA used for *in vitro* release studies was purchased from Generi BioTech, sense strand: 5’-AlexaFluor488-r(GAA GGU CGG UGU GAA CGG AU) d(TT)-3’ and antisense strand: 5’-r(AUC CGU UCA CAC CGA CCU UC) d(TT)-3’. High Pure RNA Isolation Kit and Cell Proliferation Reagent WST-1 were purchased from Roche. QIAshredder and RNeasy Fibrous Tissue Mini Kit were purchased from Qiagen. Gene-specific TBP primers: forward: 5’-ACC CTT CAC CAA TGA CTC CTA TG-3’ and reverse: 5’-TGA CTG CAG CAA ATC GCT TGG-3’ and control RNA (ctrlRNA) forward: 5’-r(CCU GCA CCA CCA ACU GCU UAG) d(AA)-3’ and reverse: 5’-r(CUA AGC AGU UGG UGG UGC AGG) d(TT)-3’were obtained from Integrated DNA Technologies IDT. TPP Petri dishes and TPP 24-well plates from BioTech. Gentamicin from Sandoz, Novartis Company. MMP-9 Monoclonal Antibody (5G3) and GAPDH Loading Control Monoclonal Antibody (GA1R) were purchased from Invitrogen. 3M™ Tegaderm™ Transparent Film Dressing was obtained from I.T.A. Interact, Tissue Adhesive was obtained from Surgibond, Glucometer FreeStyle Freedom Lite with cartridges were purchased from Abbott, and Disposable Biopsy Punch was obtained from Stiefel.

Poly(vinyl alcohol) (PVA 40-88 and PVA 5-88) was purchased from Merck and poly(L-caprolactone) from Sigma Aldrich. Chemicals (synthesis, purification and origin) for preparation of copolymer-functionalized nanodiamonds in section 2.2 were described in detail in [26].

### 2.2. Synthesis of cationic copolymer-functionalized fluorescent nanodiamonds (Cop^+^-FND)

Commercially available high pressure high temperature diamond nanocrystals (Microdiamond MSY 0–0.05 µm, type Ib with nitrogen impurity ∼200 ppm) were oxidized by air in a furnace at 510 °C for 4 h and treated with a mixture of HF and HNO_3_ (2:1) at 160°C for 2 days. The nanocrystals were washed with water, 1 M NaOH, 1 M HCl and water (for detailed centrifugal parameters see [30]). To create NV centers in the diamond lattice, we used a previously published protocol (see ESI in [31]). Briefly, the powder was irradiated with a 16.6 MeV electron beam followed by annealing (900 °C, 1 h) and air oxidation (510°C, 4 h). The powder was again treated with a mixture of HF and HNO_3_, washed with NaOH, HCl, water and then friezed–dried. The resulting powder was redispersed in Milli-Q water (60 mL, 2 mg mL^-1^) by probe sonication (for sonication parameters see [26]). Transparent colloid containing fluorescent nanodiamonds (FNDs) was incubated for 30 min at room temperature and filtered using a 0.2 µm PVDF filter. The particles were coated with a methacrylate-terminated silica layer using a modified Störber procedure[32]; the amount of all components needed for silication procedure was linearly recalculated (according to [26]) to the amount of FND powder (120 mg). The terminal methacrylate groups on the FND surface allow to proceed with a radical polymerization resulting in a dense layer of statistical cationic copolymer poly{(2-dimethyl-aminoethyl methacrylate)-*co*-[N-(2-hydroxypropyl)methacrylamide]} (poly(DMAEMA-co-HPMA)); Detailed preparation (including purification of the chemicals) and characterization of FND@silica@poly[DMAEMA-*co*-HPMA] complexes was shown in [26] (specifically, sample denoted as “80^+^/20^0”^ in [26] was used in this study). Briefly, HPMA and AIBN were freshly recrystallized prior to use. Both monomers DMAEMA (1968 mg) and HPMA (656 mg) were dissolved in DMSO (7.5 mL); AIBN (752 mg) was added to the mixture and filtered using 0.2 µm PTFE filter. Methacrylate-terminated FNDs in DMSO (120 mg, 752 µL) were added and the stirred mixture was secured in argon. The polymerization proceeded for 3 days under argon atmosphere at 55 °C. The reaction was terminated by MeOH addition; resulting Cop^+^-FND sample was purified by centrifugation using nuclease free water (sample was manipulated in laminar flow box). Final concentration after purification was 74.6 mg mL^-1^ and the sample was stored at 4 °C. A long-term storage conditions: Cop^+^-FND in MeOH at –20 °C.

### 2.3. Electrospinning of hybrid PVA/PCL nanofiber mesh containing Cop^+^-FND:siRNA / siRNA-A488 / ctrlRNA

PVA 40-88 powder was dissolved in nuclease-free water (nfH_2_O) at a concentration of 15 % (w/v) and heated for 30 min at 90 °C under stirring. After cooling to RT, the PVA solution was sonicated for 1 min (Bandelin Sonorex^TM^ RK 31, 30/240 W, 35 kHz) and autoclaved (121 °C, 15 min). Desalted siRNA lyophilizate was dissolved in nfH_2_O (500 µM, 7.3 mg mL^−1^); 32.2 µL of the siRNA stock solution was mixed with 66.8 µL of autoclaved PVA solution (15 % (w/v)). To create Cop^+^-FND:siRNA complex respecting Cop^+^-FND: siRNA mass ratio 32: 1: siRNA-PVA mixture was placed in sonication bath filled with DEPC-treated water and 101.0 µL of Cop^+^-FND (74.6 mg mL^−1^) dissolved in nfH_2_O were slowly pipetted (not dropwise) directly into the sonicated mixture of siRNA-PVA. The resulting dispersion was kept sonicating for 30 s after mixing (sample was placed in sonication hotspot). Resulting stable deep brown colloidal dispersion (200 µL) of Cop^+^-FND:siRNA in 5% (w/v) PVA solution was formed. A typical picture of this dispersion was shown in [28] (see ESI Figure S4e in [28]). This procedure was repeated ten times to obtain 2 mL of dispersion. All samples (including sonication bath) were manipulated in a laminar flow box. The same procedure was used to complex Alexa Fluor 488 labeled siRNA (siRNA-A488) and non-active siRNA (ctrlRNA).

Nanofibers were prepared using InoSPIN MINI electrospinning unit (InoCure) with needle electrode (G10) and rotating drum collector (100 mm diameter) covered with aluminum foil. The voltage in the system was set at potential difference of 45kV and temperature of 30°C. The composites were prepared by spinning of 20 ml PCL nanofibers from acetic:formic acid (7:3) solution with concentration of 20 (w/v) %. On the top of the solution, nanofibers from PVA solution containing either active or non-active siRNA were deposited. The solution of PVA with Cop^+^-FND:siRNA (2 mL) was mixed with 2.5 mL 30% PVA 5-88, 3.75 mL 20% PVA 40-88 and 6.75 mL dH_2_O with total 10 (w/v) % PVA concentration. In addition, 40,000 ppm of glyoxal (w_glyox_/w_PVA_) and 30,000 ppm of phosphoric acid (w_glyox_/w_phosphoric acid_) were added for further crosslinking of the mesh. The mesh after spinning was stored in vacuum foil and crosslinked in oven at 60 °C for 24 hours. The resulting nanofiber mesh composite, NF{Cop^+^-FND:siRNA}, had a size of approximately 10 × 10 cm, containing 24 µg siRNA/cm^2^ NF. The NF{Cop^+^-FND:siRNA} was cut into round/square pieces with an area of 0.3 and 2.0 cm^2^. Additionally, PVA/PCL nanofiber mesh without Cop^+^-FND:siRNA complexes was used as a control and prepared with parameters similar to that of the NF{Cop^+^-FND:siRNA}. All samples for *in vitro* / *in vivo* testing were sterilized in UV light for 15 minutes prior to use.

### 2.4. Cop^+^-FND:siRNA colloidal characterization evaluated by DLS and ELS

To evaluate colloidal stability of Cop^+^-FND:siRNA in 5% (w/v) PVA, dynamic (DLS) and electrophoretic (ELS) light scattering analysis was performed. Z-average diameter and apparent ζ-potential were measured using Zetasizer Nano ZSP equipped with a He–Ne laser emitting power of 10 mW at a 633 nm wavelength. Size results were inferred from the second-order time intensity autocorrelation function *g*^(2)^(τ)*-*1 considering 3^rd^ order cumulant analysis in Zetasizer Software 7.13; sample was measured three times with an automatic duration at a backscatter angle of 173°; dynamic viscosity and refractive index of the solvent at 25 °C was set at 0.887 cP and 1.33. Apparent ζ-potential was measured using phase analysis light scattering at angle of 13°. Sample was measured two times with a dip cell considering Monomodal analysis; to calculate the apparent ζ-potential value from Henry’s equation, we have used the Smoluchowski approximation for spherical noncoated particles without inspecting the value of the ratio of particle size to Debye length. For both analysis, Cop^+^-FND:siRNA in 5% (w/v) PVA was diluted by pure Milli-Q water with a final volume of 0.6 mL and Cop^+^-FND concentration of approximately 0.1 mg mL^−1^; measurements were performed in disposable cuvette and dilution of the Milli-Q water by the testing sample was lower than 1 %. Reported values represent a mean value of these measurements.

### 2.5. Fluorescence lifetime imaging microscopy (FLIM) of nanofiber mesh composites

Spatial distribution of Cop^+^-FND:siRNA complexes embedded in nanofiber mesh was characterized usingFLIM. Images were recorded using the inverted time-resolved MicroTime 200 (PicoQuant GmbH) confocal fluorescence microscope. FLIM and point measurements were performed using a 531 nm sub-nanosecond pulsed laser excitation with a repetition rate of 20 MHz and avarage power of approx. 35 μW. Nanofiber samples were placed between two round glass coverslips which were gently pushed together and imaged by Olympus 60× UPlanSApo water-immersion objective with N.A. 1.2. A single-photon avalanche photodiode was used to detect fluorescence in the 641 ± 75 nm spectral region corresponding to FND luminescence. Final image resolution was 256 × 256 pixel; pixel dwell time 2 us. Fluorescence decays and lifetime images were reconstructed and analyzed by custom-made TTTR Data Analysis software.

### 2.6. Electron microscopy (FESEM)

Sample was fixed to a holder with double-sided tape and coated with a thin layer of Au/Pd. The morphologies of samples were observed by using a Hitachi S-4700 field emission scanning electron microscope (FESEM) at 15LkV. The image analysis was perfomed using ImageJ 1.53k. All micrographs from Figure S2 were separately displayed in ImageJ using 200% magnification; image was virtually dived into five different areas (image corners and middle part) and 250 manual measurements of fiber diameter were taken from these selected areas (50 measurements/area; 250 measurements/image; 8×250 measurement/sample). The micrographs (8 images per sample) were recorded from distinct locations.

### 2.7. Release of Cop^+^-FND:siRNA from nanofiber mesh in physiological-like conditions

Release of the complexes from nanofiber mesh was assessed spectrophotometrically using Alexa Fluor 488 labelled siRNA (siRNA-A488). Round NF{Cop^+^-FND:siRNA-A488} composite possessing an area of 0.3 cm^2^ was incubated for 14 days in 1 mL of PBS in a humid atmosphere consisting of 5 % CO_2_ and 37 °C. Controls (PBS buffer and NF without nanoparticles in PBS buffer) were incubated in the same manner. At given incubation times (24, 72, 144, 216 hours), the fluorescence intensity was directly measured in the supernatant using Infinite M200 Pro plate reader (Tecan), Ex/Em 488 nm/525 nm. The maximal releasable amount of Cop^+^-FND:siRNA-A488 from the composite was determined after the total dissolving of the NF{Cop^+^-FND:siRNA-A488} composite in CHCl_3_ without further purification. The final concentrations of released components from NF{Cop^+^-FND:siRNA} can reach the following maximal values at the end of the experiment: 800 nM (i.e. 12 μg mL^−1^) siRNA and 370 μg mL^−1^ Cop^+^-FND at defined Cop^+^-FND: siRNA mass ratio 32: 1.

### 2.8. Inhibition of MMP-9 metalloproteinase expression

#### In vitro RNA isolation

NIH/3T3 CRL-1658™ cells were maintained at 37 °C in a humidified atmosphere containing 5 % CO_2_ in plastic Petri dishes. The growth medium contained DMEM, 10 % FBS and 44 mg L^−1^ of Gentamicin. Fresh medium was supplied three times a week. To ensure the exponential growth phase of the cells, they were passaged twice a week. For testing of silencing effectivity of NF{Cop^+^-FND:siRNA}, NIH/3T3 cells were seeded in concentration 125,000 cells well^−1^ in 1 mL of DMEM supplemented with 10% FBS in plastic 24-well plates. Cells were left for 4 hours to adjust and adhere to the bottom. Round NF{Cop^+^-FND:siRNA} composite with the area of 2.0 cm^2^ was added to the media on top of the cells and cells were incubated for further 144 hours at 37 °C, 5 % CO_2_. Cells without the addition of NF{Cop^+^-FND:siRNA} were used as a control. Cell lysis and total RNA extraction were performed using a High Pure RNA Isolation Kit.

#### Scar tissues RNA isolation

RNA from tissues was extracted using cell homogenizer QIAshredder and purified using RNeasy Fibrous Tissue Mini Kit.

#### RT-qPCR assay and analysis

Isolated RNA (1 µg per sample) from NIH/3T3 CRL-1658™ cells and scar tissues were transcribed to cDNA using High-Capacity cDNA Reverse Transcription Kit. MMP-9 mRNA level was detected with TaqMan Universal Master Mix and MMP-9 specific PCR primer. TBP housekeeping gene was used as an internal control. It was detected with Power SYBR™ Green PCR Master Mix and primers: forward 5’-ACC CTT CAC CAA TGA CTC CTA TG-3’and reverse 5’-TGA CTG CAG CAA ATC GCTTGG-3’. Samples were analyzed in technical triplicates. CFX96 Touch Real-Time PCR Detection System (Bio-Rad) was used to measure probes fluorescence and CFX Manager 3.1 software (Bio-Rad) was used for sample analyses. The expression of the MMP-9 was normalized to the expression of TBP and fold change was calculated by the software (based on 2^-ddCt^ method). Expression of MMP9 and TBP mRNAs in tissues were tested after the formation of primary scars (days 35-42).

### 2.9. Cytocompatibility of NF{Cop^+^-FND:siRNA} composite

NIH/3T3 cell line was seeded in 96-well plastic plates, 20,000 cells well^−1^ in 200 μL of DMEM containing 1 % FBS. NF{Cop^+^-FND:siRNA} composite (round area of 0.3 cm^2^) was placed on top of the cells and incubated for 48 hours in 37 °C, 5% CO_2_.

#### LDH cytotoxicity assay

At the end of the incubation time, supernatant (100 µL) was mixed with 100 µL of reagent mixture and incubated for 30 minutes. 50 µL of stop buffer was added and absorbance at 490 nm was measured with the Infinite M200 Pro plate reader (Tecan) using iControl software, reference measurement at 630 nm. Lysis buffer from the kit was used as a positive control.

### 2.9 Induction of diabetes-like conditions in animal models

Diabetic-like conditions were established by the administration of low dose Streptozotocin (STZ) injection to 8-10 weeks old C57BL/6 mice (*Mus musculus,* male) according to ref. [33]. Animals’ procedures followed the Czech law regarding animal protection and the Etical Committee of the Czech Academy of Sciences approved the experimental plan with numbers 36/2015 and 80/2017. Glucose levels were measured with Abbott glucometer after 6 hours of starving on day 0 of the experiment. STZ was dissolved in sodium citrate (pH 4.5) immediately before use and it was administered intraperitoneally for five consecutive days. The daily dose comprised 1 mg of STZ per 25 g of animal weight. Animals were starved for 4 hours before each STZ injection. On day 19, glucose level was measured to see its difference from day 1. We reached 90 % of animals with elevated glucose levels defined as: glucose level >8.3 mmol/l after 6 hours of starving. These animals were considered to mimic diabetic-like conditions. Prior wounding (day 22), the animals were anaesthetized using ketamine-xylazine cocktail (100 µL/20 g of animal weight; containing 0.16 mg of xylazine and 1 mg of ketamine) and they were shaved. Wounds were created using Disposable Biopsy Punch of 0.6 cm in diameter (0.3 cm^2^). The tissue taken during wounding was kept in RNAlater^TM^ stabilization solution for further analysis. Wounds were covered with nanofiber mesh (0.3 cm^2^ sized squares) and moistened with 50 µL of PBS. Transparent film dressing (Tegaderm^TM^) was then used to protect the wound site and it was fixed with tissue adhesive (Surgibond) placed on the non-wounded part of the skin. Wounds analysis were taken on the wounding day (day 22), on day 29 and days when primary scars were formed. The wounds were considered healed when the primary scars were formed and scar tissues were excised and stored in RNAlater^TM^ for further analyses.

### 2.10 Activity and amount of MMP-9 protein in healing and scar tissues

Excised tissues were lysed with M-PER™ Mammalian Protein Extraction Reagent. Total protein concentrations in lysates were measured by Pierce™ Coomassie Plus™ (Bradford) Assay Kit. Bovine serum albumin standard of known concentrations (from 0–10 µg µL^−1^) was used to define a standard curve. Seven µL of standard or sample were mixed with 193 µL of Bradford reagent and pipetted in the transparent 96-well plate. Plate was incubated for 10 minutes and the absorbance at 595 nm was measured using an Infinite M200 Pro plate reader (Tecan). The protein concentrations in samples were determined from the standard curve.

MMP-9 enzymatic activity was assessed using zymography. Protein containing sample (5 μg of protein per 5 µL) was mixed with 5 µL of Novex Tris-Glycine SDS Sample Buffer. Electrophoresis was executed in an XCell SureLock^TM^ Mini-Cell using Novex Running Buffer and following settings: running time 90 minutes, constant voltage 125 V, and starting current 40 mA. Novex Sharp Pre-Stained Protein Standard served for band identification. Renaturing Buffer was used to wash the gel after electrophoresis for 30 minutes at room temperature. The gel was then incubated in Developing Buffer for 90 minutes. Bands on the gel were visualized by incubation with SimplyBlue SafeStain for 1 hour, followed by 1-hour incubation in dH_2_O.

The MMP-9 protein level in wound was assessed by western blotting and compared to GAPDH protein load control. Total 160 µg of protein sample (adjusted up to 16.3 µL with dH2O) were mixed with 2.5 µL of Reducing Agent and 6.3 µL of LDS Sample. The sample mixture (25 µL) was loaded onto the 10-well NuPAGETM 4-12% Bis-Tris Mini gel. Novex Sharp Pre-Stained Protein Standard was loaded to determine the molecular weight and transfer efficiency. Electrophoresis ran for 35 minutes at constant 200 V using XCell SureLockTM Mini-Cell and NuPAGETM MES 1x Running Buffer. Samples were then transferred onto the PVDF membrane using a semi-dry blotter and settings: 90 minutes, constant voltage 7 V, and starting current 30 mA. The membrane was washed in 0.1 % Tween-20/PBS (T-PBS), blocked in 5% non-fat milk for 1 hour, and incubated with 1 mL of primary antibody MMP-9 Monoclonal Antibody (5G3) overnight (1:1000 antibody in T-PBS). Next day, the membrane was washed in 0.1 % T-PBS and incubated in 10 mL of secondary antibody for 60 minutes (1:200 antibody in T-PBS; Goat Anti-Mouse IgG Peroxidase Conjugated). After washing in 0.1 % T-PBS, the membrane was incubated with SuperSignal™ West Pico PLUS Chemiluminescent Substrate. A G: Box Chemi XRQ and GeneSys software (both Syngene) were used for band detection and analyses. GAPDH control was detected accordingly using primary antibody GAPDH Loading Control Monoclonal Antibody (GA1R).

### 2.11 Statistical analysis

Survival analysis of the time-to-event data (time to wound closure) was performed by the Weibull regression model in the Bayesian framework (Figure 5) using software packages in R [34–41]. Probability density function rW((), underlying survival sW(() and hazard hW(() functions of a Weibull random variable *r* were given by Equations (1)–(3):

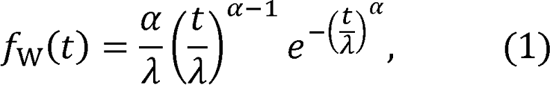

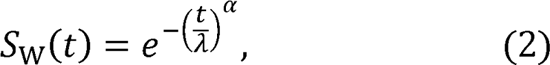

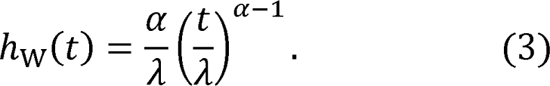

The Bayesian model defined below was fitted with **brms** package [39] that employs Stan software for probabilistic sampling.

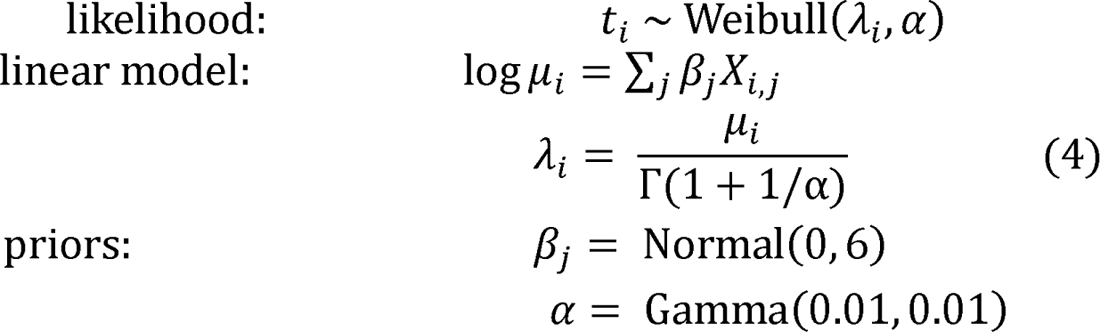

The first row shows a stochastic part of the model and states that the response variable *r* is a random variable independently drawn from a Weibull distribution with shape parameter *a* and scale parameter 1. The linear model in the second line describes how logµ_i_ (log of the conditional mean for the response variable) is constructed for given treatment,; x_i,j_ is an indicator variable. The remaining lines describe weakly informative prior distributions. Unlike **brms** Weibull likelihood, the linear model is parametrized in terms of log_µi_, the back-transformation of estimated p_j_ parameters into 1 metric was performed by Equation (4) [42]; Γ is the gamma function. Markov chain Monte Carlo sampling procedure was specified by four chains with 5000 iterations per chain, of which 2000 serve as a warm-up. Model diagnostic indicates convergence of the sampling chains of the parameters (trace rank plot visual check, w = 1.00, number of effective samples >6700). The full computer code including more detailed model diagnostics and criticism (prior sensitivity analysis, posterior predictive check etc.) is provided in R Markdown document [43]. The graphical output was tailored using Inkscape graphics editor [44].

**Figure 1.**
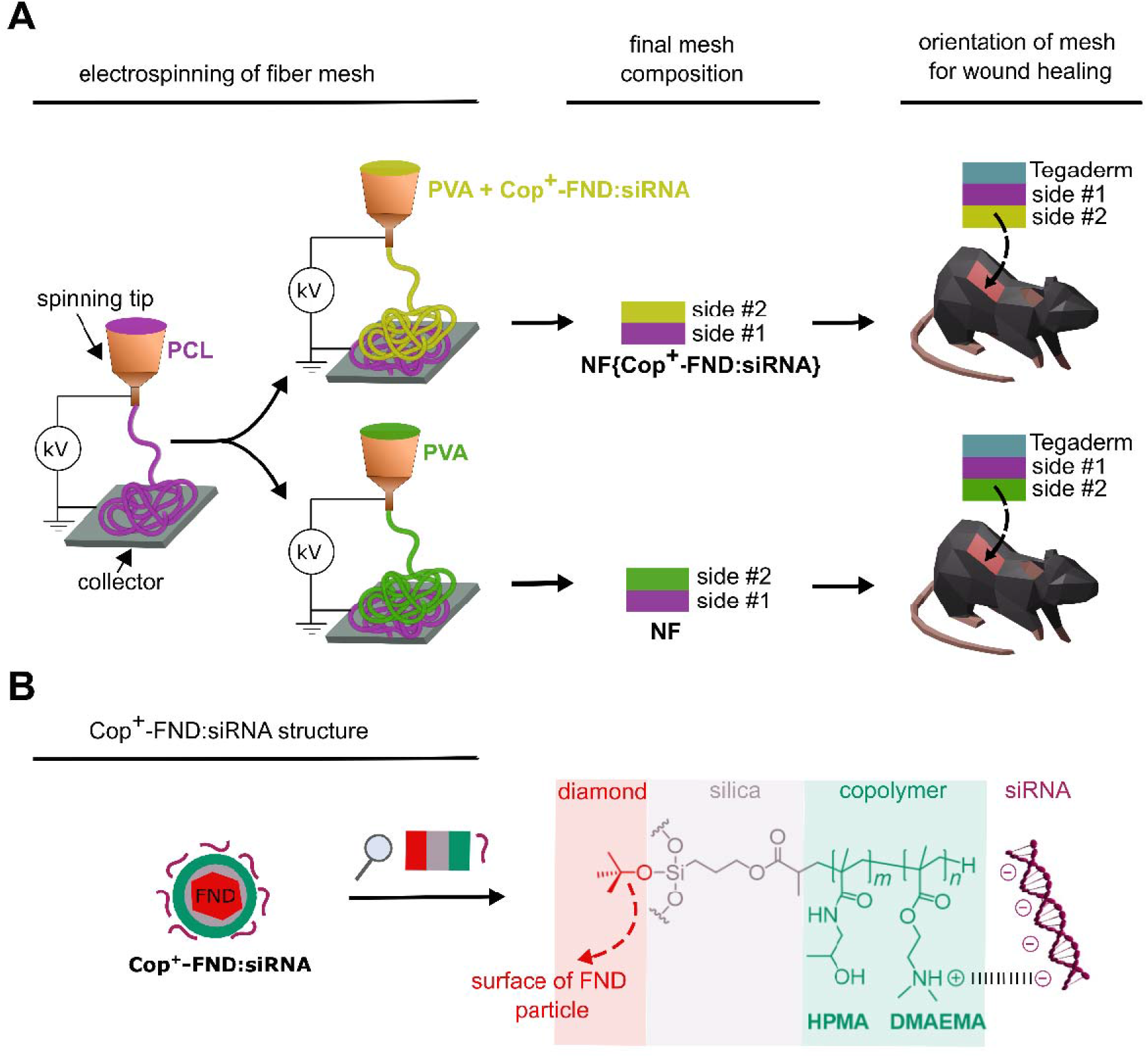
**(A)** Preparation procedure of hybrid PVA/PCL fibrous mesh containing Cop^+^-FND:siRNA complexes *–* NF{Cop^+^-FND:siRNA} (upper part), and control mesh without nanodiamond complexes *–* NF (bottom part). Side #2 is facing the wound during the treatment. **(B)** Schematic structure of cationic copolymer coating on the surface of fluorescent nanodiamond (Cop^+^-FND), which binds electrostatically the siRNA molecule.

**Scheme 1.**
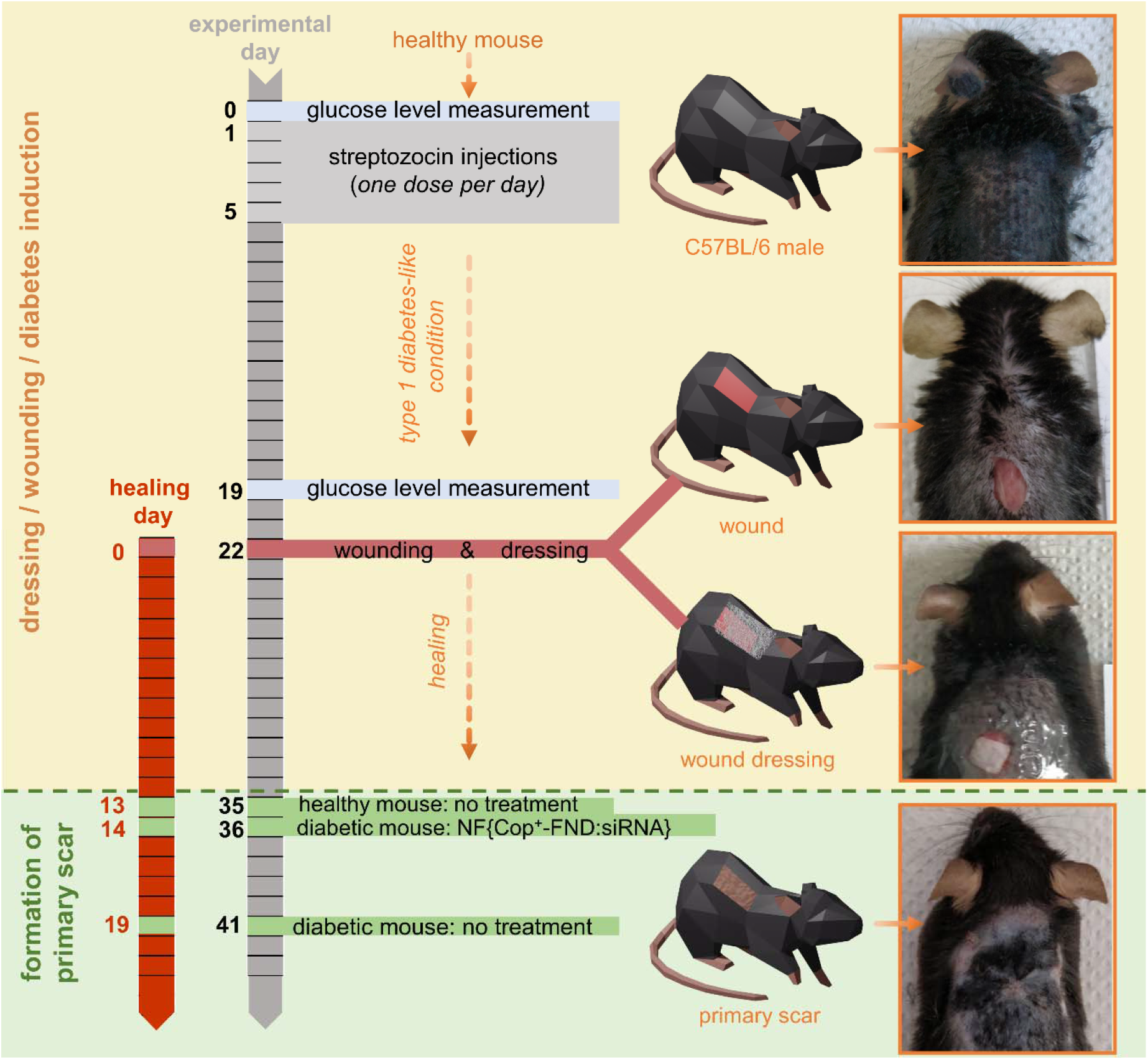
Time schedule of wound healing model preparation. **Orange top area:** induction of diabetes-like conditions in healthy C57BL/6 mice model (days 0–22) followed by wound creation and application of primary (nanofiber-based) and secondary (Tegaderm) dressing (day 22). **Green bottom area:** the wounds were considered healed when the primary scars were formed. The experiment was performed independently three times.

## 3 Results and discussion

### 3.1 The NF{Cop^+^-FND:siRNA} composite – preparation and characterization

In the recent studies, we focused on the colloidal robustness [26], intracellular fate [27] and biodistribution study of Cop^+^-FND:siRNA complexes [28]. We observed that the localized intratumoral application of the colloidally stable Cop^+^-FND:siRNA led to high inhibition efficacy in mice bearing xenografted Ewing sarcoma tumor.

Here, we utilized the potential of this delivery system for localized *in vivo* therapy and took advantage of its combination with hybrid biodegradable PVA/PCL nanofiber mesh designed for topical wound healing. Unmodified nanodiamonds have been successfully incorporated in nanofibers of various compositions [45–47]. We designed and fabricated by electrospinning a nanofiber mesh composite NF{Cop^+^-FND:siRNA} which has two main functions: primary bioactive dressing and a scaffold for controlled release of Cop^+^-FND:siRNA complexes into the wound.

First, we focused on estimation of the optimal Cop^+^-FND:siRNA mass ratio. An optimal mass ratio should ensure (i) negligible free fraction of siRNA after complexation with Cop^+^-FND, (ii) positive apparent ζ-potential of the complexes after siRNA binding, (iii) mass ratio as low as possible to reduce the amount of Cop^+^-FNDs in downstream applications, but (iv) still keeping good colloidal stability of the system. In good agreement with the previous studies [27], we chose the Cop^+^-FND:siRNA mass ratio 32:1, which fulfills the above listed criteria (i)-(iv). Moreover, the powerful electro-sterical stabilization of Cop^+^-FND:siRNA complexes provided by the copolymer coating allows direct complexation of Cop^+^-FND with siRNA even at high particle concentrations such as 75 mg mL^−1^ without formation of aggregates (see **Figure 3A**). Thus, highly concentrated/low-volume samples can be easily produced, which is a typical requirement for *in vivo* applications. Apparent Z-average diameter of the Cop^+^-FND:siRNA complexes in presence of the main dispersing component of the nanofibers, PVA, was 91.3 ± 1.5 nm (measured by DLS, see **Figure 3A**). However, one should be aware that a free diffusion of the complexes is likely restricted due to extension of the electrical double layers surrounding the particles in low ionic strength solvent as Milli-Q water. The magnitude of a cumulant based polydispersity index (PDI) 0.12 ± 0.02 represents a lower limit of mid-range PDI values and the sample can be considered as nearly monodisperse. The relative measure of a part of the double-layer charge, which also contributes to the interparticle interaction, was characterized by apparent ζ-potential 43.3 ± 2.2 mV (measured by ELS). All values represent mean ± standard deviation, a more detailed information can be found in **Table S1** in Supporting Information.

Next, we incorporated the Cop^+^-FND:siRNA complex in the hybrid PVA/PCL nanofiber mesh by blend electrospinning, providing NF{Cop^+^-FND:siRNA}. We utilized a combination of highly hydrophilic polyvinyl alcohol homopolymer and slowly biodegradable [48] poly-lJ-caprolactone homopolyester provides a balanced nanofiber for wound healing. As was shown in [49], PCL and PVA are highly suitable for formation of nanofibrous wound dressings. In the current setting, the PCL fibers were used as integrative part of wound dressing stimulating adhesion and proliferation of skin cells. The PCL fibers were spun as the first layer subsequently covered by PVA mesh containing the bioactive nanoparticles. PVA was used as a hydrophilic polymer electrospun from aqueous solutions and enabling dispersion of Cop^+^-FND:siRNA nanoparticles. The controlled release of PVA was regulated by crosslinking of PVA chains which alternates nanofiber degradation kinetics [50]. We determined the morphology of the electrospun scaffold containing the NF{Cop^+^-FND:siRNA} using field emission scanning electron microscopy (FESEM). PVA/PCL mesh without the particles served as a control. The resulting FESEM micrographs revealed the fibrous morphology of both samples (Figure 2, left column; **Figure S2**). However, the side #2 of NF{Cop^+^-FND:siRNA} sample showed a higher content of microfibrous fraction in comparison with side #2 of nanoparticle free control (see Figure 2B, 2C and S2). The different morphology could be explained by the rheological behaviour of highly concentrated nanoparticle dispersion stabilized by a linear polyelectrolyte (Cop^+^-FND). Similar colloidal systems are typically influenced by electroviscous effects (e.g. swelling of linear chains), which are caused by overlapping of the electrical double layers in the first approximation. Consequently, an increase in solution viscosity and surface tension can be observed. These parameters belong among the most important and affect the fiber diameters as well as the resulting mesh morphology.

**Figure 2.**
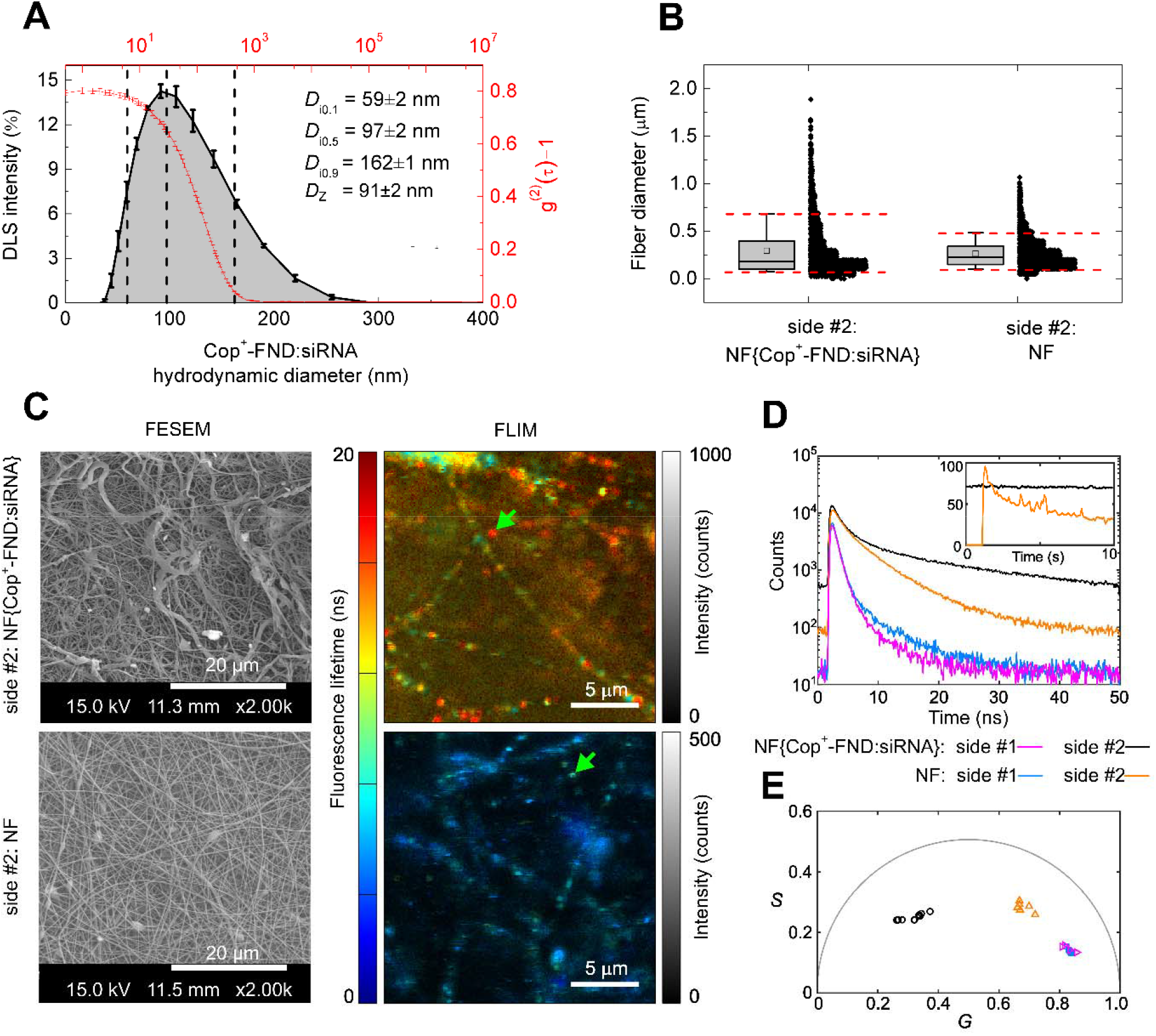
(**A**) DLS measurement of Cop^+^-FND:siRNA in 5% PVA (sample was further diluted by Milli-Q water according to the section 2.4). Intensity size distribution was inferred from the raw intensity autocorrelation function *g*^(2)^(τ) –1. Vertical black dashed lines coupled with the histogram represent particle diameters *D*_i0.1_, *D*_i0.5_ and *D*_i0.9_ at 10^th^, 50^th^ and 90^th^ percentiles of the size distribution; *D*_Z_ represents Z-average diameter. The error bars represent standard deviations over three single measurements from separately prepared samples. (**B**) Fiber diameter quantification (box plot and all measured data points organized in histogram bins) obtained from image analysis of FESEM micrographs (for all analyzed images see **Figure S2**). Box plots characterize a fiber diameter distribution using 25^th^, 50^th^ and 75^th^ percentiles, and 90% and 10% whiskers; the horizontal line in the box: median of the distribution; square: mean of the distribution. Number of fibres analysed per sample was 2000. (**C**) Comparison of nanofibers NF{Cop^+^-FND:siRNA} with a control nanofiber sheet without particles (NF). Only side #2 facing the wound is shown for both samples – NF{Cop^+^-FND:siRNA} (top images) and NF (bottom images). **Left column:** FESEM micrographs – for all images see **Figure S2**; **middle column:** FLIM images with green arrows indicating locations of single-point measurement. (**D**) Fluorescence decays acquired for 10 s from a single point from both sides of NF{Cop^+^-FND:siRNA} and NF, as indicated in the legend. Inset shows intensity traces of the measurements from side #2. (**E**) A phasor plot of all recorded fluorescence decays – each point corresponds to a FLIM image or point-measurement.

To assess the distribution of Cop^+^-FND:siRNA in NF{Cop^+^-FND:siRNA} composite, we investigated the particles using FLIM. The long emission lifetime of fluorescent nitrogen-vacancy centers in FND (approximately 25 ns; [51]) enabled us to distinguish the Cop^+^-FND:siRNA complexes from the short-living autofluorescence background of the nanofibers (Figure 2C, FLIM). The presence of Cop^+^-FND:siRNA complexes in the side #2 of NF{Cop^+^-FND:siRNA} was unambiguously confirmed by both FLIM images recorded from several areas and single-point measurements from both sides of NF{Cop^+^-FND:siRNA} and NF. Exemplary fluorescence decays are shown in the Figure 2D. Furthermore, photobleaching was observed during acquisition from side #2 of NF while no decrease of signal was observed either during imaging or single-point measurements from NF{Cop^+^-FND:siRNA} as can be expected for photo-stable luminescence of nitrogen-vacancy centers. Intensity traces during 10 s single-point measurement are shown in the inset of Figure 2D. To represent the rather complex fluorescence decays, we employed the graphical phasor approach. Figure 2E shows a phasor plot of all time-resolved data recorded from NF{Cop^+^-FND:siRNA} and NF. Each point in the plot corresponds to one measurement, i.e., either FLIM image, or a single-point decay. Clustering of the points in the phasor plot demonstrate the consistency of the time-resolved measurements and that FLIM can reliably reveal the presence of long-lived NF{Cop^+^-FND:siRNA}. Further, as the FLIM images were recorded from several distinct locations and various scanned area sizes (ranging from 20×20 to 80×80 µm), we can conclude that the Cop^+^-FND:siRNA complexes can be found in abundance and are distributed homogenously in the NF{ Cop^+^-FND:siRNA} composite.

In summary, we confirmed the intended structural arrangement of the sample suitable for wound dressing and controlled release of Cop^+^-FND:siRNA in the wound. Most importantly, we found a high loading of Cop^+^-FND:siRNA on the side #2 of NF{Cop^+^-FND:siRNA} (facing the wound). In contrast, we did not detect the Cop^+^-FND:siRNA complex either on the side #1 of NF{Cop^+^-FND:siRNA} or on any side of the NF control sample.

### 3.2 Cell culture – NF{Cop^+^-FND:siRNA} release, inhibition efficacy and cytotoxicity

For the effective topical treatment, a controlled release of the Cop^+^-FND:siRNA complexes from the hybrid NF{Cop^+^-FND:siRNA} mesh to the wound is essential. In a simple release experiment in PBS (Figure 3A) we found a gradual liberation of approximately 25 % of all embedded complexes within 144 h. Interestingly, this hybrid structure produced by blend electrospinning did not show the undesirable initial burst release, which is typical for scaffolds manufactured this way. We then measured the inhibition efficacy of the siRNA against MMP-9 mRNA liberated from NF{Cop^+^-FND:siRNA} composite in NIH/3T3 CRL-1658 cell culture. After 144 h incubation with the NF{Cop^+^-FND:siRNA}, MMP-9 mRNA was extracted and its fraction was measured compared to reference by RT-qPCR. Figure 3B shows that NF{Cop^+^-FND:siRNA} treatment reduced the MMP-9 mRNA expression to 17 ± 33% (mean ± standard deviation) indicating a specific inhibition compared to the treatment with NF control. To assess a potential cytotoxicity of the tested scaffolds, we monitored the release of lactate dehydrogenase from damaged cells (Figure 3C). We did not observe any adverse effects after 48 h.

**Figure 3.**
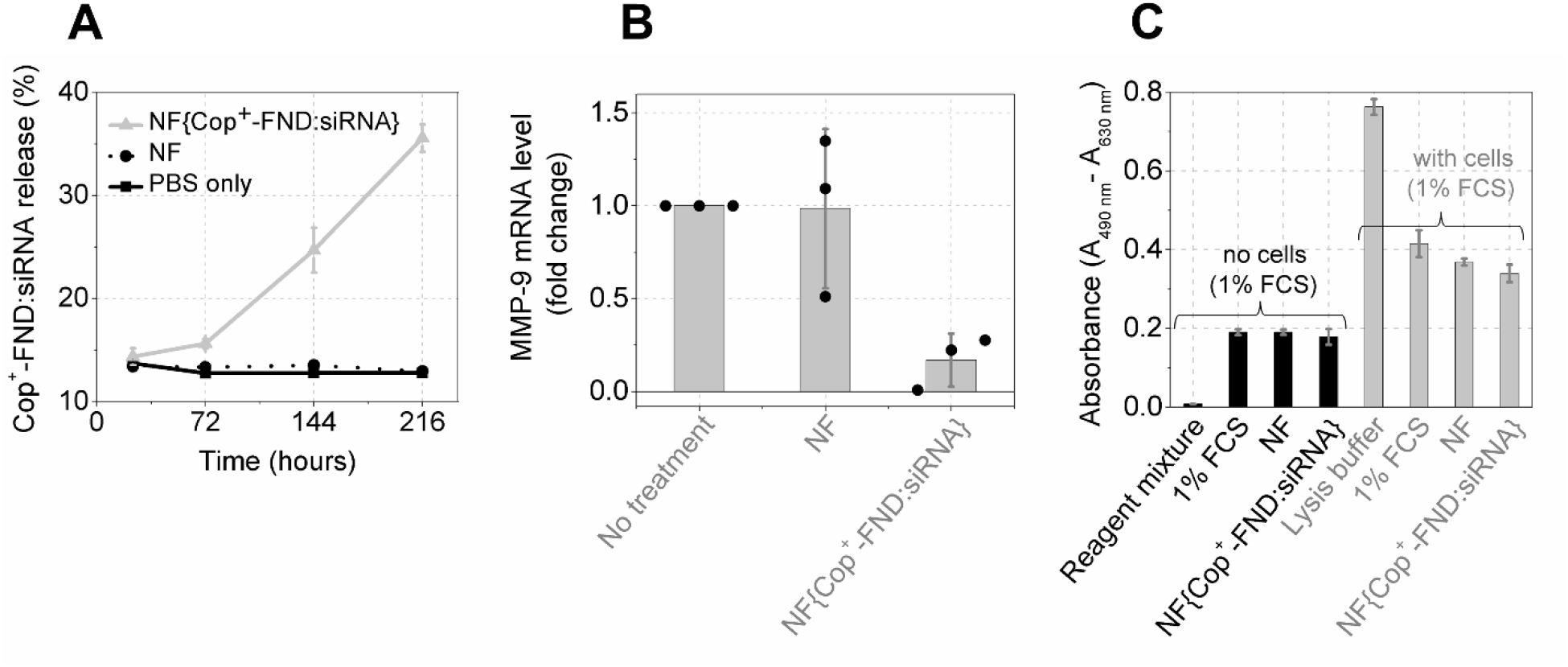
(**A**) Time-dependent release of Cop^+^-FND:siRNA from nanofiber mesh in physiological-like conditions (PBS buffer, pH = 7.4, 37 °C). Control samples: NF – nanofiber mesh without nanoparticles, 1x PBS without additives. (**B**) *In vitro* inhibition of MMP-9 mRNA expression in NIH/3T3 CRL-1658 cells in culture using nanofiber mesh (2.0 cm^2^ mL^-1^) determined 144 h post-treatment by RT-qPCR. (**C**) Cell cytotoxicity: LDH assay assessing the release of lactate dehydrogenase from damaged cells 48 h post-treatment. The nanofiber mesh of area 0.3 cm^2^ / 200 μL was used, which roughly corresponds to 300 nM (i.e. 5 μg mL^−1^ siRNA) and 200 μg mL^−1^ Cop^+^-FND at defined Cop^+^-FND: siRNA mass ratio 32: 1 (≈ 15 % of all embedded complexes). Data for each analysis ((A), (B) resp. (C)) were obtained from single experiment where the samples were analysed in triplicates **–** the error bars represent standard deviations over these replicates.

### 3.3 Diabetic-like animal model – in vivo NF{Cop^+^-FND:siRNA} application

To evaluate the in vivo efficacy of our delivery system, we used a diabetic-like mouse model of chronic wound healing. This model system involves surface application enabling direct visual observation of changes triggered by used NF{Cop^+^-FND:siRNA} composite. Importantly, the improved wound healing here relies on re-establishing of disease-impaired balance and it can be achieved by targeting one solely metalloproteinase (such as MMP-9) thus representing an easy model for gene therapy.

The diabetic-like animals developed high glucose levels two weeks (experimental day 19) after STZ injections (see Figure 5C) in agreement with previously published protocol [33]. Dorsal wounds were then created (experimental day 22 / healing day 0) and subjected to wound healing treatment with NF{Cop^+^-FND:siRNA} composite and controls. To distinguish the healing efficacy after application of our NF{Cop^+-^FND:siRNA} composite, we observed the wound till a primary scar was formed (Figure 4).

**Figure 4.**
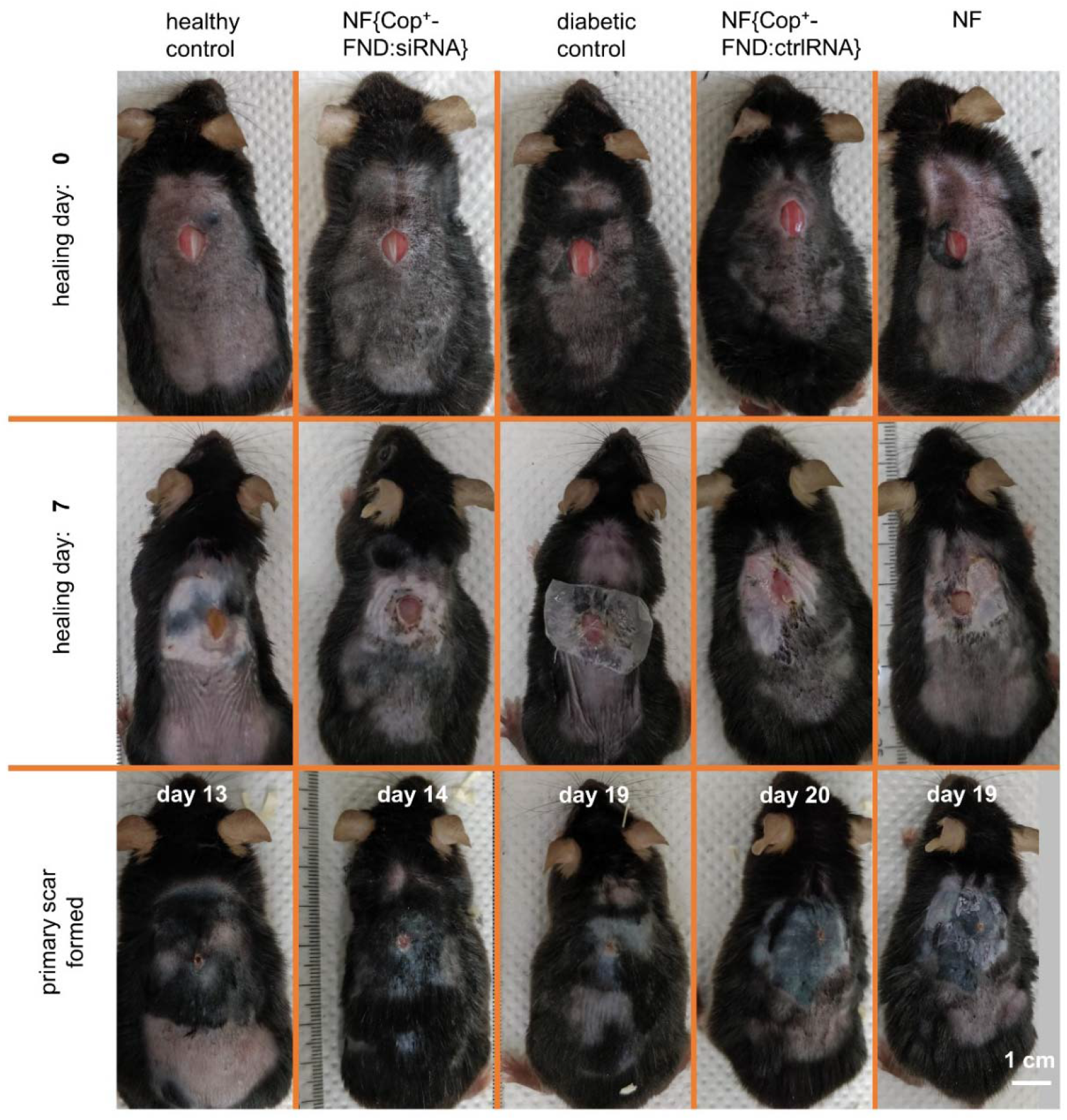
Photographic representation of diabetic-like wound healing showing different rates of primary scar formation in dependence on the type of treatment. Healthy and diabetic controls were treated only by the Tegaderm film without the primary nanofiber dressing. The rest of the mice were treated by the nanofiber dressing, which was covered by the Tegaderm film (secondary dressing). The experiment was performed independently three times (15 animals in total; 5 animals per experiment). The photographs represent the first experimental group (Group #1 – 4 diabetic-like animals and 1 healthy control).

To quantify the treatment effect size, we chose Bayesian inference as a main statistic tool since it remains valid even for very limited experimental datasets. The power/weakness of this approach is dependence upon the prior knowledge that is an inherent part of the Bayesian procedure. In other words, to obtain the estimation of the treatment effect size, we need to first specify prior information about it (based on pilot data, practical experiences etc.). This prior information is then computationally updated in light of observed data to reach the desired effect size (posterior effect size). [52,53]

Analysis of the time-to-event data (Figure 5B, C) requires techniques for positive-valued random variables (survival models). As a first-choice model, we utilized the semiparametric *Cox proportional hazards model* to compare wound closure rates. This approach makes no specific assumptions about the hazard function h(() which describes the frequency at which the wound closure occurs per unit of time, given that the event hasn’t happened yet (instantaneous wound closure rate). The obtained results (not shown) were strongly dependent on the choice of the prior information, and thus, not very useful for further discussion. Imposing more strict assumptions on the hazard function leads to fully parametric models [54]. The basic single-parameter exponential model states that the hazard function is constant over time. This is not a very reasonable expectation for the wound closure process. This expectation would imply that a nonnegligible fraction of animals would experience wound closure after a very short time after the wounding (e.g. after two days). Two-parameter models bring more flexibility when concerning hazard profiles. Aware of this, we utilized the two-parameter Weibull model which allows us to assume a more feasible monotonically increasing hazard function hW(() (see Equation 3 for a> 1) [42,53]. In other words, the chance of wound closure increases with time. Results of the Weibull regression model in Figure 5A, B (smooth curves) are shown in terms of the survival function s_W_(() (see Equation 2) which describes the probability that the wound closure has not occurred by elapsed time.

The unique feature of the Bayesian inference is to provide not only one estimate of the survival function for a given treatment but rather a set of survival functions with different relative plausibilities (Figure 5A, B – the shaded area surrounding the dashed curve could be graphically replaced by a set of survival curves for given treatment). Thus, one can estimate e.g. a set of times until 50% of mice experienced the wound closure for given treatment, resp. set of median survival times “*T50”;* (sW(() = 0.5). Figure 5D shows the relative plausibilities for the obtained *T50* times with shaded areas under the curves that indicate an arbitrary range of *T50* times compatible with the data and utilized model (89% compatibility interval) [52]. In terms of *T50* (means of *T50* distributions in Figure 5D), healing of the wounds in healthy animals took 12.9 days. As expected, the scar formation was delayed in diabetes-like conditions by 6.7 days.

We found that the application of our NF{Cop^+^-FND:siRNA} composite promoted healing efficacy by 5.7 days in comparison with diabetic control without treatment. Thus, the wounds in diabetic-like animals were healed 29 % faster than in diabetic-like animals without the fibrous bioactive dressing. The healing efficacy of NF{Cop^+^-FND:siRNA} treated diabetic-like animals was then similar to healthy animals without diabetes-like conditions (delayed by 1.0 day).

Taking into account the overall position of the posteriors in Figure 5D for NF and NF{Cop^+^-FND:ctrlRNA} relative to diabetic-like control, the results are most compatible with no important effect. In contrast, posteriors for healthy control and NF{Cop^+^-FND:siRNA} are considerably separated from the control samples. All these results bring convincing preclinical evidence about the effectivity of NF{Cop^+^-FND:siRNA} treatment.

**Figure 5.**
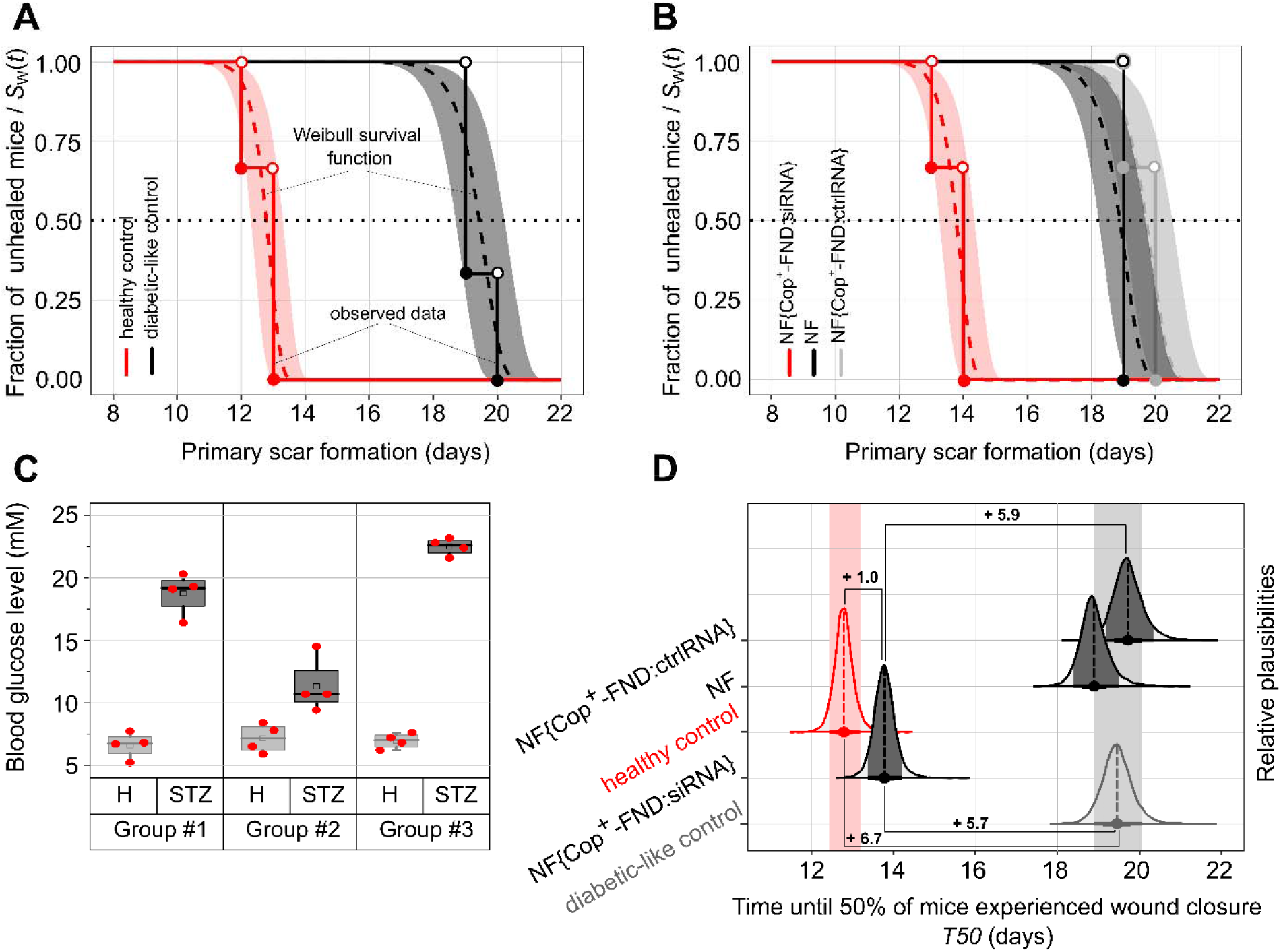
(**A**) Healing of healthy and diabetic-like controls without treatment and (**B**) comparison of -like animals treated with nanofiber-based composites. Step functions show measured data; time to the formation of the primary scar (*n* = 3 per group). Dashed curves show the survival functions estimated by Weibull regression in the Bayesian framework; shaded areas surrounding the dashed curves represent 89% compatibility intervals; see section 3.3 for more detailed explanation. (**C**) Blood glucose level of healthy animals (experimental day 0; H = “healthy”) and animals after induction of diabetes-like conditions (experimental day 19; STZ = “Streptozotocin”) – three independent groups (*n* = 4 per group) were tested; Box plot description is the same as for Figure 2B**. (D)** Posterior relative plausibilities of median survival times *T50* obtained from estimated survival functions; shaded areas under the posterior curves show 89% compatibility intervals; dashed vertical lines represent a mean of the posterior distribution. Results represent data from three independent experimental groups (15 animals in total). The *x*-axis for (**A**) and (**B**) reflects healing days as indicated in **Scheme 1**.

Regarding the model limitations, the applied two-parametric form of the Weibull regression model assumes not only the monotonic profile of the hazard functions but also their proportionality. In other words, the survival curves (e.g. in Figure 5A, B) are not allowed to cross among treatments [55]. In addition, one could alternatively consider the application of the three-parameter Weibull model (a third shift parameter is included) due to the moderately large Weibull shape parameter (a= 38) obtained by the two-parametric version [56].

Relative MMP-9 mRNA expression measured by RT-qPCR (Figure 6A) showed a 54 ± 10 % decrease when comparing scar tissue exposed to the NF{Cop^+^-FND:siRNA} (experimental days 35–42) and tissue before treatment (experimental day 22). Further comparison of relative MMP-9 mRNA downregulations between NF{Cop^+^-FND:siRNA} and NF{Cop^+^-FND:ctrlRNA} reveals 2.5 ± 0.4-fold decrease (54 ± 10%: 22 ± 1%) of MMP-9 expression (Figure 6A). A similar result was achieved for the diabetic control.

**Figure 6.**
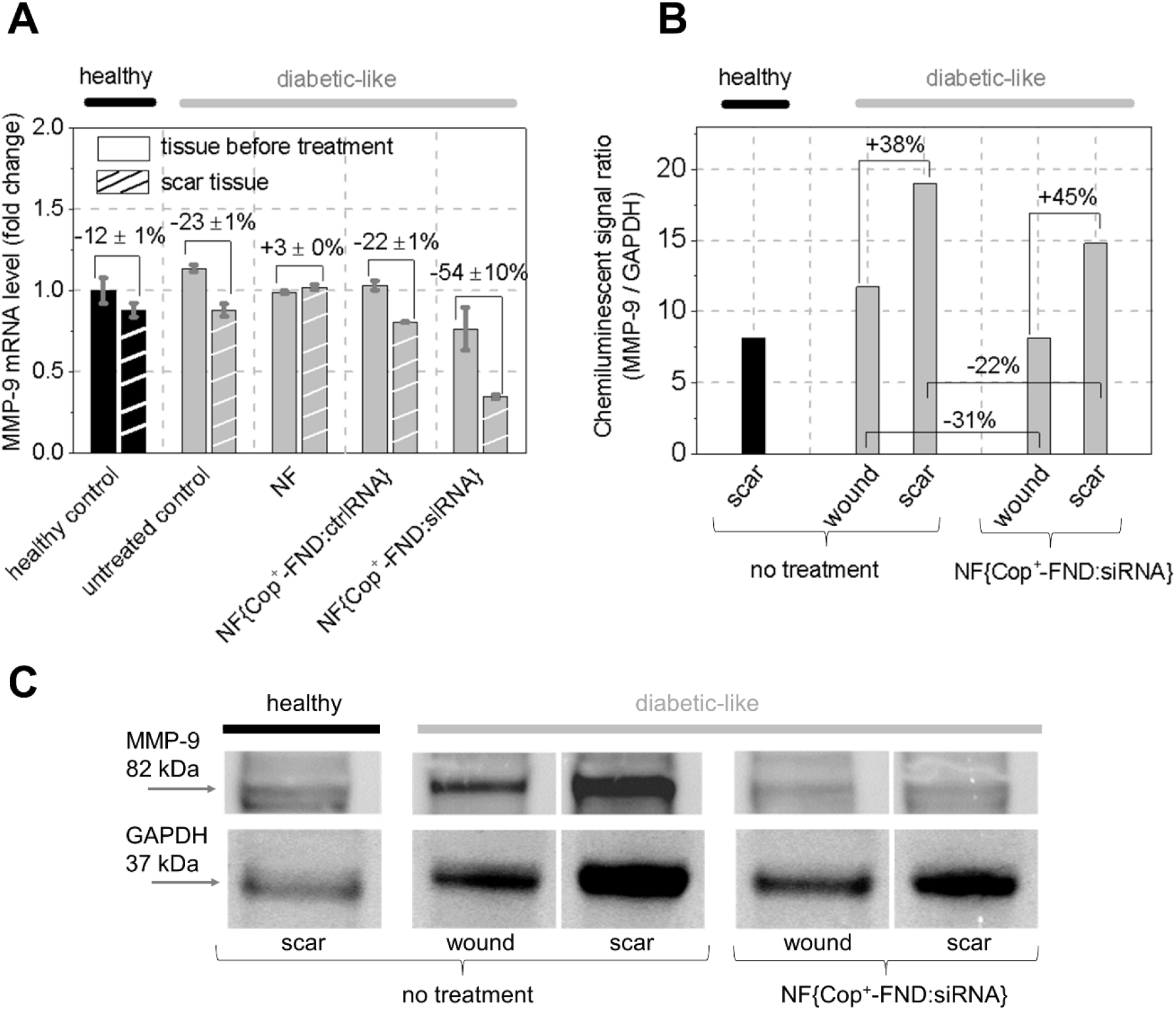
**(A)** Expression of MMP-9 mRNA in tissues before treatment (experimental day 22) and in scar tissues (experimental days 35–42) excised from healthy and diabetic-like animals. **(B)** Level of MMP-9 protein in excised tissues after 7 days of treatment (experimental day 29, samples denoted as wound) and after scar formation (denoted as scar) in healthy and diabetic-like tissues analyzed densitometrically from western blot **(C)**.

The MMP-9, enzyme responsible for wound ECM degradation, is secreted by neutrophils, fibroblasts, macrophages, and endothelial cells in the form of precursor pre-MMP-9. Pre-MMP-9 is cleaved by matrix metalloproteinase 3 resulting in its catalytic domain exposure and maturation. [57]

We observed specific activity of MMP-9 proteinases in healing wound and scar tissues (see zymogram in **Figure S1**). The activity of MMP-9 was reduced in animals treated with NF{Cop^+^-FND:siRNA} composite. To quantify the amount of MMP-9 protein we used western blot analysis followed by band densitometry evaluation (Figure 6B, C). The amount of MMP-9 protein was increasing during the healing progression towards scar formation as the protein accumulated in the wound site. Using NF{Cop^+^-FND:siRNA} we decreased the amount of MMP-9 protein during the course of healing and scar formation by 31 % and 22 %, respectively, relative to the non-treated control (Figure 6B).

Overall, our findings support the unique benefit of gene therapy by restoring MMP-9 balance in damaged tissue while keeping resident cells without observable adverse effects. In a critical view, the used artificial wound model in diabetic-like mice represents a relatively rough approximation of a human chronical wound. A typical disadvantage of this model is a significant wound contraction observed in the normal process of mice wound healing, while bottom-up epithelialization and granulation represent normal wound healing in human [58]. However, application of nanofiber and in lesser extent also Tegaderm dressing improves quality of a scar tissue [59]. Due to moisture retention, the nanofiber-based dressings promote re-epithelialization and reduce epidermal thickness resulting thus in good imitation of human wound healing [60]. It is reasonable to expect that MMP-9 imbalance also affects the wound contraction [58]. On the other hand, the high similarity level between murine and human MMP-9 and the possibility to incorporate fully degradable nanoparticles (instead of Cop^+^-FND) inside the fibrous mesh [61] would allow easier translation of this pre-clinical model into reasonable clinical research.

## 4 Conclusion

In summary, we designed hybrid biodegradable PVA/PCL fibrous mesh embedded with previously established cationic copolymer-grafted fluorescent nanodiamonds carrying siRNA targeted against MMP-9 mRNA (Cop^+^-FND:siRNA). We assembled the hybrid nanomaterial using blend electrospinning and characterized its structure by FLIM and SEM. The fibrous mesh showed a controlled release of the Cop^+^-FND:siRNA complexes without an undesirable initial burst release. The bioactive dressing allowed conservation and sustained release of the siRNA in the murine diabetic-like wound. We have demonstrated delayed wound healing in diabetic-like animals compared to animals without diabetic conditions due to MMP-9 deregulation. Biofunctionality of the fibrous mesh containing Cop^+^-FND:siRNA complexes was demonstrated by a significant reduction of the healing time – scar formation for treated diabetic-like animals became comparable with non-treated diabetes-free mice. Our healing system emphasizes the importance of selective gene therapy which may circumvent the common, non-specific treatments of topical chronic injuries such as non-healing diabetic ulcers. The restoration of MMP-9 balance in damaged tissue seems to be a promising way which may ease the translation of topical gene therapy into the clinic.

## Acknowledgement

The authors are grateful to Dr. David Chvatil (Institute of Nuclear Physics of the CAS) for the irradiation of nanodiamonds by electrons and to Dr. Zuzana Zlamalova Cilova (University of Chemistry and Technology, Prague) for her support with FESEM measurements. This work was supported by Czech Academy of Sciences – Strategy AV21 (VP29), and by European Regional Development Fund; OP RDE; Project: CARAT (No. CZ.02.1.01/0.0/0.0/16_026/0008382).

## Supporting Information

**Figure S1.**
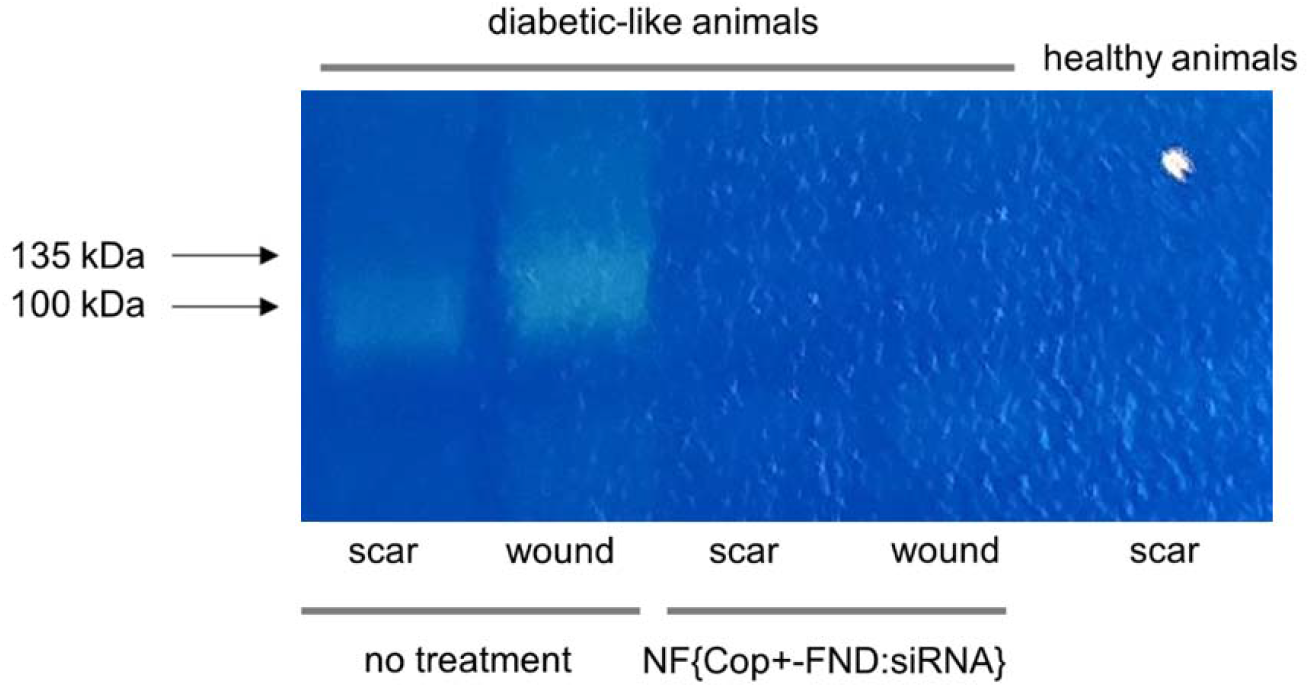
Activity of pre-MMP-9 and MMP-9 in wound tissues after 7 days of treatment (wound; experimental day 29) in diabetic-like tissues revealed by zymography. Scar tissues from diabetic and healthy animals excised in the end of experiment are shown for comparison (scar). Precursor MMP-9 and matured MMP-9 migrated at 135 kDa and 100 kDa, respectively.

**Figure S2.**
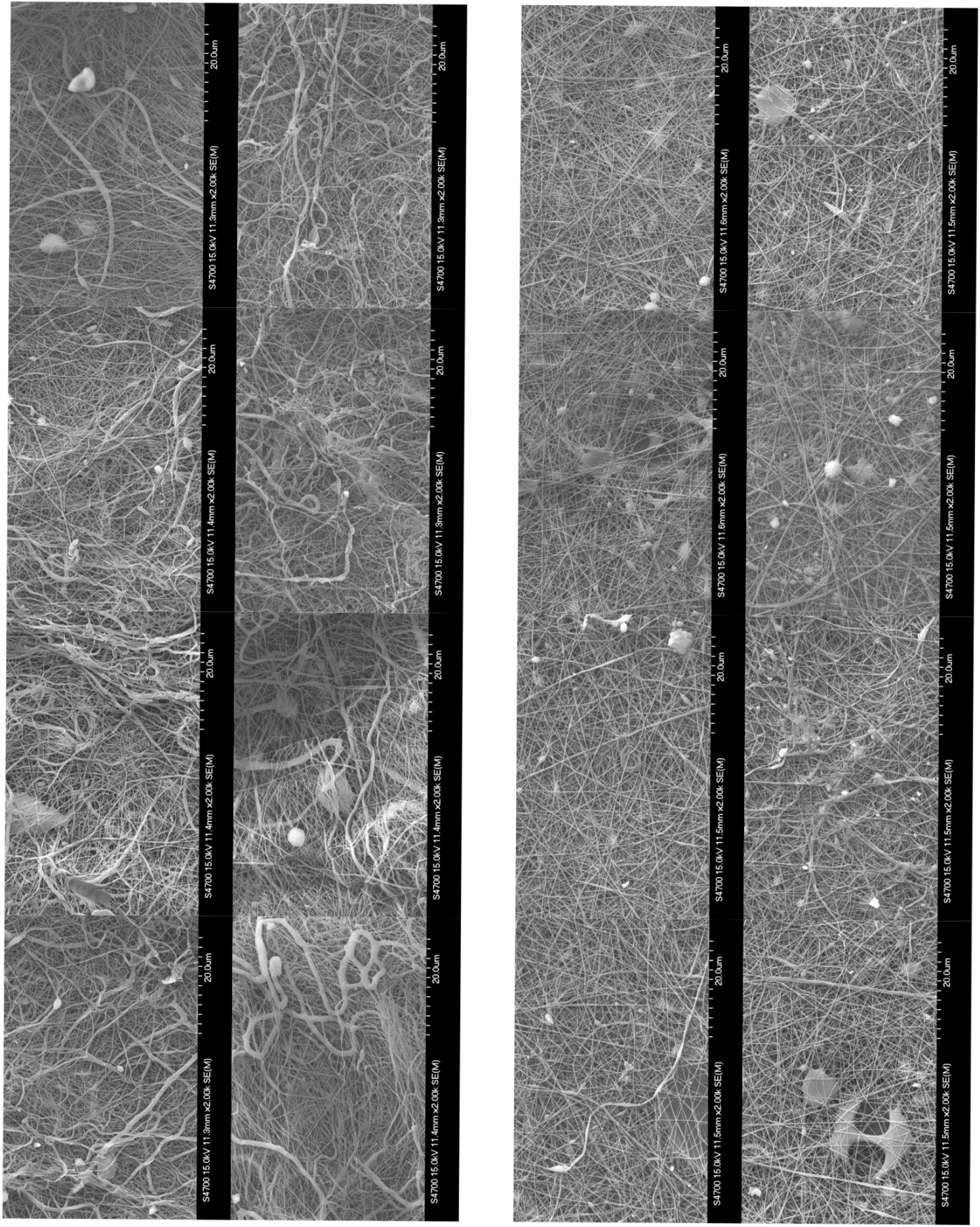
FESEM micrographs (only side #2) used for image analysis – **Figure 2B** – **left column:** NF{Cop^+^ FND:siRNA}; **right column**: NF.

---

**Table S1.**
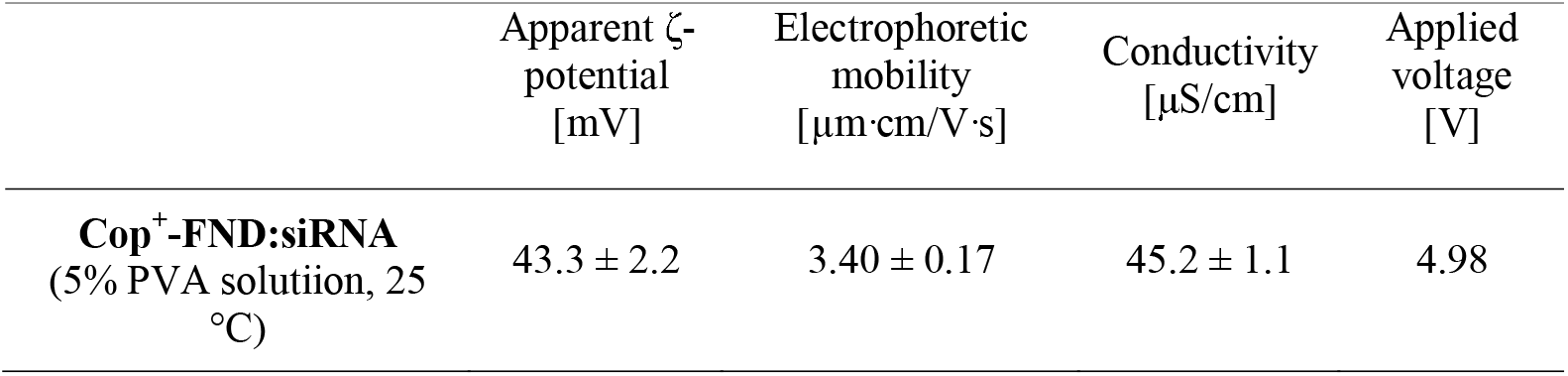
Electrophoretic light scattering analysis of the Cop^+^-FND:siRNA complexes (mass ratio 32: 1).

| | Apparent ζ-potential [mV] | Electrophoretic mobility [$\mu\text{m}\cdot\text{cm}/\text{V}\cdot\text{s}$] | Conductivity [$\mu\text{S}/\text{cm}$] | Applied voltage [V] |
| --- | --- | --- | --- | --- |
| **Cop<sup>+</sup>-FND:siRNA**<br>(5% PVA solution, 25 °C) | $43.3 \pm 2.2$ | $3.40 \pm 0.17$ | $45.2 \pm 1.1$ | 4.98 |

